# A spatial statistical framework for the parametric study of fiber networks: application to fibronectin deposition by normal and activated fibroblasts

**DOI:** 10.1101/2022.06.14.496046

**Authors:** Anca-Ioana Grapa, Georgios Efthymiou, Ellen Van Obberghen-Schilling, Laure Blanc-Féraud, Xavier Descombes

## Abstract

Changes in the spatial landscape of the extracellular matrix (ECM) in health and disease significantly impact the surrounding tissues. Quantifying the spatial variations in the fibrillar architecture of major ECM proteins could enable a profound understanding of the link between tissue structure and function. We propose a method to capture relevant ECM features using graph networks for fiber representation in normal and tumor-like states of 4 alternatively spliced isoforms of fibronectin (FN) associated with embryonic development and disease. Then, we construct graph-derived statistical parametric maps, to study the differences across variants in normal and tumor-like architectures. This novel statistical analysis approach, inspired from the analysis of functional magnetic resonance imaging (fMRI) images, provides an appropriate framework for measuring and detecting local variations of meaningful matrix parameters. We show that parametric maps representing fiber length and pore orientation isotropy can be studied within the proposed framework to differentiate among various tissue states. Such tools can potentially lead to a better understanding of dynamic matrix networks within the tumor microenvironment and contribute to the development of better imaging modalities for monitoring their remodeling and normalization following therapeutic intervention.

**Author Summary:** Due to the complex architectural diversity of biological networks, there is an increasing need to complement statistical analyses with a qualitative and local description. The extracellular matrix (ECM) is one such network for which fiber arrangement has a major impact on tissue structure and function. Thus, a flexible numerical representation of fibrillar networks is needed for accurate analysis and meaningful statistical comparison of ECM in healthy and diseased tissue. First, we propose a versatile computational pipeline to study fiber-specific features of the ECM with graph networks. Then, we introduce a novel framework for the statistical analysis of graph-derived parametric maps, inspired from the statistical analysis of fMRI parametric maps. This analysis is useful for the quantitative/qualitative comparison of ECM fiber networks observed in normal and tumor-like, or fibrotic states. These methods are applied to study networks of fibronectin (FN), a provisional ECM component that dictates the organization of matrix structure. From 2D confocal images we analyzed architectural variations among 4 alternatively spliced isoforms of FN, termed oncofetal FN, that are prevalent in diseased tissue. We show how our approach can be used for the computation and statistical comparison of heterogeneous parametric maps representing FN variant-specific topological/geometrical features. These methods may be further developed and implemented into tumor tissue ECM profiling to decipher the specific roles of ECM landscapes and their remodeling in disease.

## Introduction

Organs and tissues are composed of a non-cellular stromal compartment known as the extracellular matrix (ECM), a cell-derived bioregulatory scaffold that provides chemical and mechanical cues to surrounding cells. The composition and architecture of the ECM is tissue- and organ-specific, and depends on the pathophysiological state of the tissue (i.e. normal vs diseased) (1). For example, while healthy interstitial tissues display a loose meshwork-like ECM, their fibrotic and cancerous counterparts are characterized by the presence of dense aligned ECM fibrils. Thus, the physical and structural traits of the tumor matrix have recently drawn much attention as cancer hallmarks and potential therapeutic targets (2)(3). Collagen, among the most prominent matrix components, has been extensively investigated in this context and several studies addressing its structural features and their association to cancer progression, metastasis and treatment have been published (4)(5)(6). Collagen deposition, however, depends on another key component of the ECM, namely fibronectin (FN). FN is a dimeric glycoprotein that forms a provisional matrix framework to which other ECM components integrate to generate a mature ECM (7)(8).

FN exists in two distinct forms: plasma FN (pFN) that is constitutively produced by the liver and circulates at high concentrations in blood plasma, and cellular FN (cFN) which is expressed and deposited locally in tissue by specialized cells in a spatially and temporally restricted manner. Indeed, cFN is highly expressed during embryogenesis and in certain physiological and pathological conditions including inflammation, wound healing, and cancer. Structurally, the two forms differ by the presence in cFN of alternatively spliced sequences of about 90 amino acids, termed Extra Domain B (EDB) and Extra Domain A (EDA), that enhance FN assembly, impact matrix architecture, and tune cellular responses (9).

Despite the pivotal role of FN and its variants in health and disease (10), comprehensive studies of FN fiber features have not been reported. To that end, we set out to develop a robust pipeline of numerical analyses for the extraction of biologically relevant metrics from confocal microscopy images of FN variant-specific matrices. First, to capture and study the relevant ECM fibrillar features, we developed a graph-based approach that recapitulates the fiber networks. Starting from these graph representations of FN variants, we then designed a framework to statistically compare ECM geometrical features using different parametric maps, such as fiber length and pore orientation isotropy. Using these approaches, we show here that significant parametric differences across variants in health and tumor-like states can be quantified and simultaneously localized.

## Results and Discussion

### Generation of FN variant-specific cell-derived matrices and induction of a tumor-like state

Fibroblasts are the major ECM-producing cells of tissues. During tumor progression, quiescent fibroblasts become activated by environmental cues that induce their phenotypic conversion from a “normal” to a “pro-tumoral” state (Fig 1A, top). This process is characterized by actin reorganization and increased contractility of cells that result in their elongation and the deposition of a highly anisotropic ECM (9). In vitro, these changes can be mimicked by treating normal fibroblasts with Transforming Growth Factor-β (TGF-β), a potent cytokine involved in fibroblast activation in the tumor microenvironment (9) (Fig 1A, bottom). For our analyses, normal or tumor-like FN-variant specific matrices were generated by presenting FN-null mouse embryo fibroblasts with recombinant FN containing one, both (or none) of the Extra Domains (prepared as previously described (9)), in presence or absence of TGF-β, as schematized in Fig 1B. Cell cultures were decellularized after 7 days, and matrices were visualized by immunofluorescence staining and confocal microscopy. The organization of variant-specific FN matrices in normal and activated states (Fig 1C) was then quantitatively analyzed, as described below.

**Fig 1.**
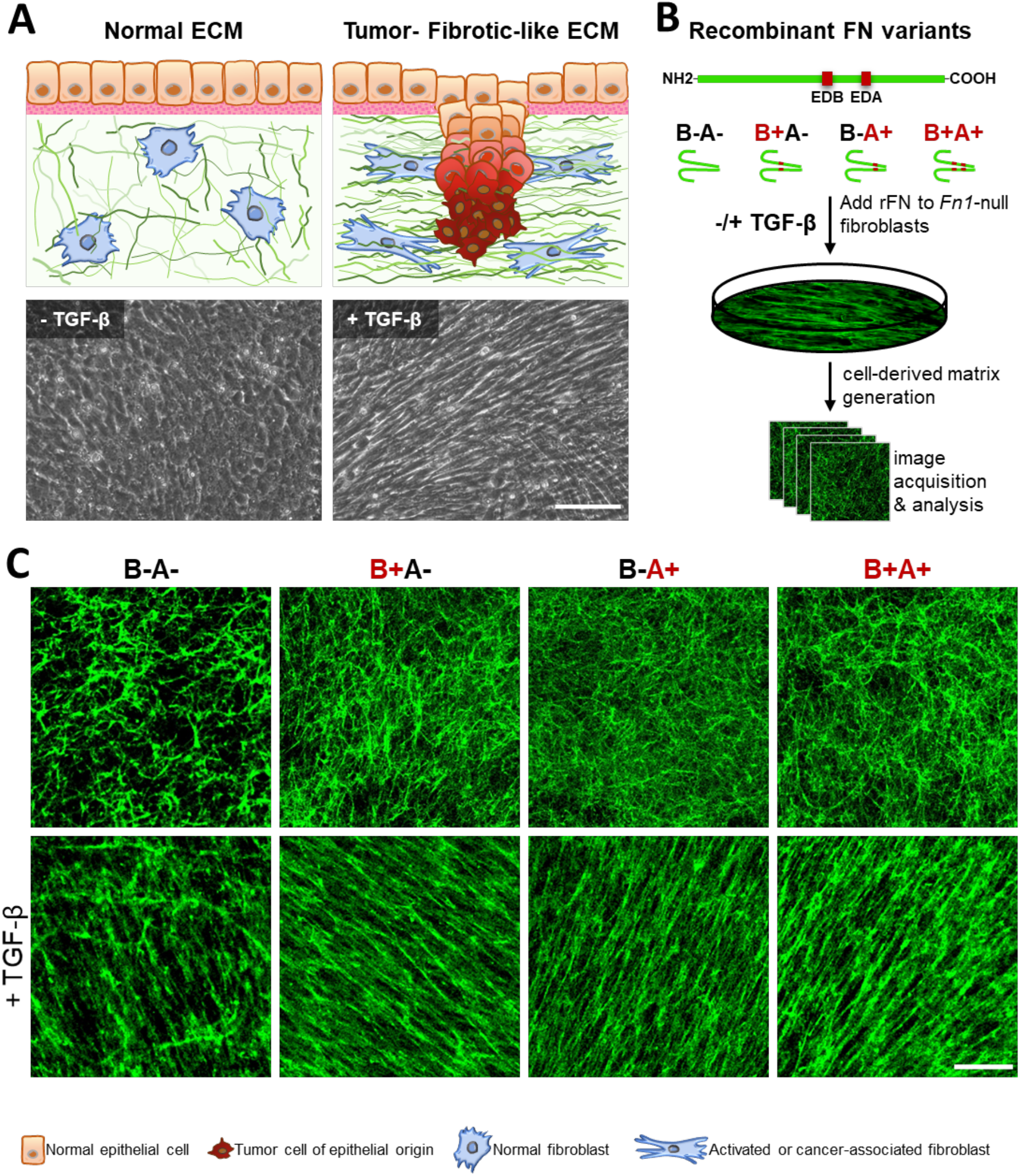
(**A**) (top row) Schematic representation of an epithelium displaying the different architectures of underlying fibroblasts and their ECM in normal and tumor settings (bottom row). Phase contrast images of cultured FN1-null mouse fibroblasts treated as indicated with 5 ng/ml TGF-β. Scale bar, 100 μm. (**B**) Workflow diagram featuring the linear structure of the purified recombinant FN variants and relative positions of the alternatively spliced Extra Domains (EDB, EDA), the generation of cell-derived matrices, and image acquisition and analysis. (**C**) Immunofluorescence staining of FN in variant-specific cell-derived matrices. FN-null mouse fibroblasts were presented with the FN variants (15 μg/ml) in the presence or absence of TGF-β (5 ng/ml). After removal of the cells, matrices were stained with a rabbit-anti-FN monoclonal antibody and visualized with confocal microscopy. Scale bar: 50 μm.

### Fiber detection using Gabor filters and graph extraction

Fiber detection for pattern quantification in FN networks was produced using a pipeline, briefly described in a previous study (1), where it was primarily utilized for the extraction of local topological fiber properties. In the present study, the workflow was extended to enable a statistical analysis of parametric maps, focusing on a global spatial statistical analysis, which is relevant for quantitative and qualitative fiber analysis across multiple scales. The computational pipeline can be tested with a graphical user interface (GUI) on single samples [e.g., immunofluorescence (IF) coupled with confocal microscopy, immunohistochemistry (IHC)], or a batch of images, the output of which can be written into .png image files and fiber specific graph/Gabor-derived features collected in .csv files. The MATLAB source code and sample images can be found on GitHub, at github.com/aigrapa/ECM-fiber-graph. The steps involved in producing a 2D graph-based fiber representation from confocal microscopy images are presented in Fig 2. First, to detect and enhance the fibrillar structures from the 2D confocal images, we selected a bank of Gabor filters (2) tailored to capture fiber elements that appear at multiple frequencies and orientations. The intrinsic parameters of a Gabor filter (S1 Text) are directly linked to corresponding physical properties (i.e. fiber thickness, fiber orientation). As a result of the filter bank convolution, the intensity of each pixel of the enhanced fibers corresponds to the Gabor filter with the highest coefficient (Fig 2). This flexible, yet meaningful detection property is an important asset over alternative multi-resolution techniques commonly used for curvilinear object detection, such as discrete curvelets (3,4), whose complex discrete implementation makes them prone to translation and rotation issues (e.g. energy dissipation in neighboring sectors in the frequency domain). In subsequent steps of the graph-based representation pipeline, a series of image morphological transformations were applied after fiber enhancement, including binarization using hysteresis thresholding and fiber morphological skeleton computation. Converting the binary fiber structure into a morphological skeleton is useful to generate a simplified representation that preserves the topological properties of the original region and allows a successive skeleton-graph association.

**Fig 2.**
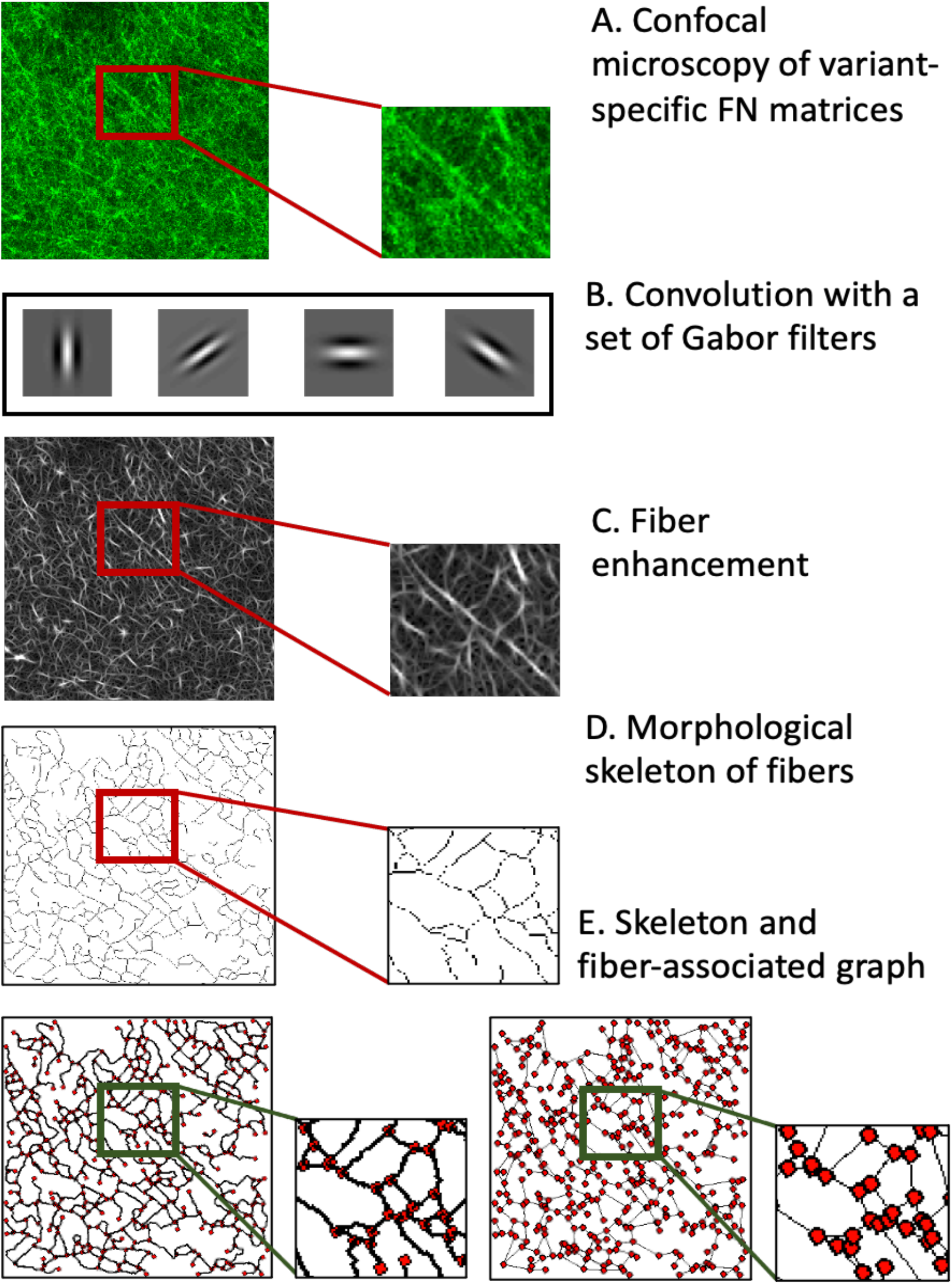
Fiber enhancement and graph-based representation starting from confocal 2D images. (**A)** Image size is set at 1024×1024 pixels, confocal acquisition is performed at a resolution of 0.27 μm/pixel. (**B)** Fibers are enhanced with Gabor filters and the maximum response is kept in every pixel, as shown in (**C**) The morphological fiber skeleton is subsequently extracted (**D**) and a fiber-associated graph is assigned (**E**). Furthermore, a simplified graph representation is derived from the skeleton graph.

Graphs (i.e. collections of nodes connected by edges) are powerful tools for the structural and pattern analysis of objects, commonly employed for the mathematical study of relations between entities (5). In our work, we propose to use a graph-based framework to depict morphological fiber skeletons, and to provide a geometrical characterization of fiber patterns. Other studies employ graphs to analyze different biological networks, such as osteocyte networks (6) and retinal vasculature (7), illustrating the remarkable potential of spatial graphs for fiber-like object analysis. Within the graph-based skeleton representation, the nodes represent either fiber crosslinks (perceived in the 2D projection of the network onto the image plane) or fiber ends, while the edges capture the fiber length between two given nodes. An additional process of node reconnection to correct for missing fibers due to artefacts from previous morphological operations and noise in the data was deemed necessary. The steps to achieve this correction are detailed in (S1 Text). As previously mentioned, the current pipeline has shown potential for the description of distinct local features among the four FN images of variants, in normal state (1).

An additional step was added to the graph-based workflow to further simplify fiber delineation. Starting from the skeleton graph, we kept all nodes corresponding to fiber extremities, and connected all pairs of nodes with a straight line, if a fiber had previously been identified. For the sake of simplicity, we refer to fiber length as the length of any straight line connecting a pair of graph nodes. This characterization can be encoded in adjacencies matrices (indicating the presence/absence) of a certain edge between the nodes. We note that both representations are useful to extract different local or global fiber properties. The graph-based skeleton fiber delineation (Fig 2) faithfully represents (according to a visual assessment performed by a trained biologist) the geometrical and topological properties of the fibers from the 2D confocal images, while the Gabor-specific (e.g. fiber local orientation, thickness) and graph-derived parameters (e.g. fiber length, number of nodes, etc.) are linked to meaningful physical fiber attributes. This biologically relevant representation enabled us to develop a statistical analysis of the variation of certain fiber parameters for both the “normal” and a “tumor-like” state of the FN network.

### Statistical analysis of parametric maps

The graph-based representation of FN networks enabled the subsequent design of a novel framework to perform a spatial statistical analysis of ECM patterns, using graph-derived statistical parametric maps. Fiber attributes computed from graphs can be considered for creating these variation maps. To illustrate an application of our method, we considered two different types of parametric maps reflecting meaningful geometrical features, namely the individual fiber lengths (i.e. the length of the connecting line between two graph nodes), and the fiber pore (“gap”) orientation isotropy (i.e. the inverse value of the absolute difference between the median and individual pore orientation). We relied on a local computation of parameters to build these maps: fiber length along graph edges or fiber gap pore orientation, and subsequent extrapolation of these values to build dense maps. Fig 3 illustrates an example for each map category. Details of the computation of parametric maps starting from graph-based representations can be found in (S1 Text).

**Fig 3.**
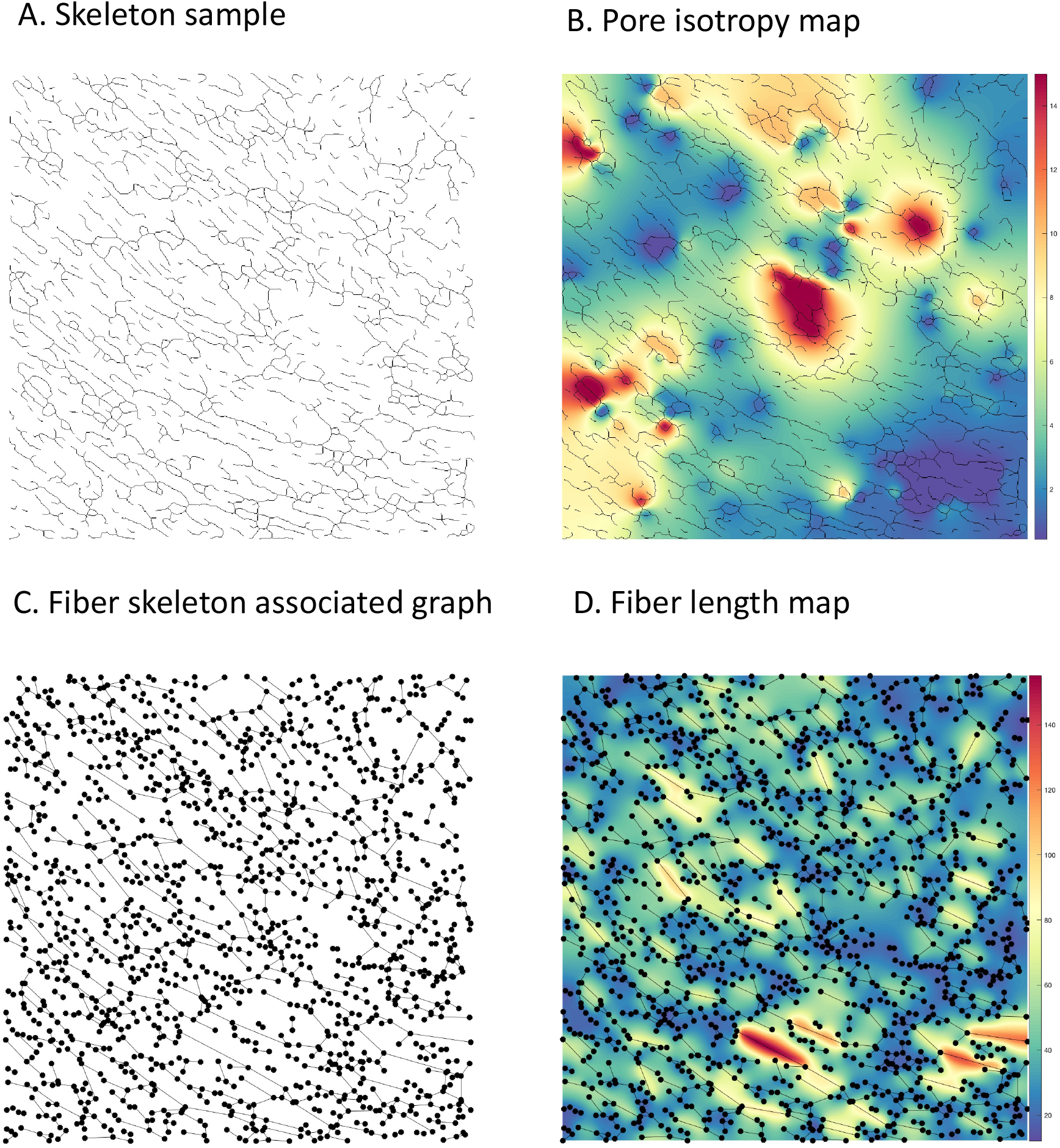
Computation of fiber parametric maps: (**A**) Starting from the skeleton graph (FN B-A+ tumor-like sample, 1024×1024 pixels), a pore isotropy map is derived (**B**), as the inverse value of the difference between the median pore angle and each individual one. (**C**) Starting from the graph-based fiber representation, a parametric map (fiber length, (**D**)) associates the fiber length, in pixels, to each corresponding connecting line.

Inspired from the statistical parameter map (SPM) framework in fMRI analysis (8,9) using Gaussian Random Fields (GRF) (10), the proposed spatial statistical learning framework aims to quantify and simultaneously localize statistically significant differences across the normal and tumor-like FN network maps. These differences are assessed both quantitatively and qualitatively, as detected anomalies with respect to the GRF, corresponding to regions within the maps that cannot be explained by the normal model. Providing both a quantitative and a qualitative assessment of the parametric variations is extremely important for the study of spatial heterogeneity of fibers in pathological conditions. Here we apply our proposed framework to determine if fiber length and pore orientation isotropy can discriminate between normal and tumoral states, and among the distinct cFN variant networks. While topographical differences between ECM of healthy and tumor tissue have been described (4), no current computational study can, to our knowledge, simultaneously localize and quantify them.

The principle of the current methodology (Fig 4) relies on modelling the normal FN maps as realizations of a GRF within a hypothesis testing framework (12) (11). We hypothesized that the tumor-like maps are realizations of the same GRF, and we determined a set of probabilities that characterize a degree of belonging to the GRF, of certain contiguous regions (clusters) at various intensity thresholds. In other words, the current statistical analysis identifies those foreign regions with respect to the GRF, within both types of maps, under the null hypothesis (i.e., at a given p-value (pval)), to compare their properties (e.g., number and size).

**Fig 4.**
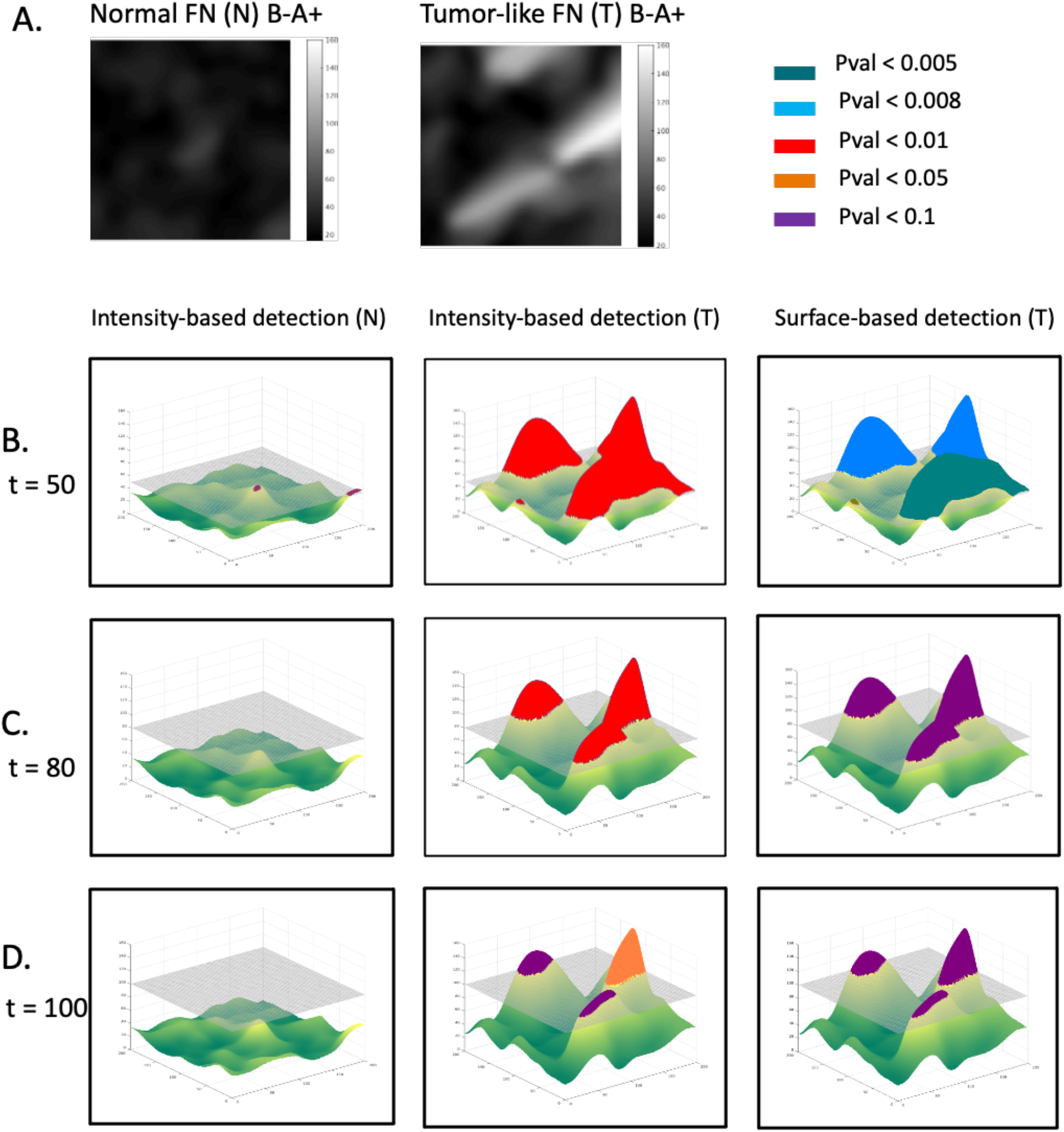
Methodology for statistical detection of foreign regions to a GRF, in normal and tumoral parametric maps. (**A**) Normal and Tumoral fiber length map (FN B-A+). The normal sample is modelled as a realization of a GRF, and we assume that the tumor-like sample is a realization of the same process. Clusters of regions with an intensity higher than a given threshold, t= 50 (**B**), t = 80 (**C**), t = 100 (**D**) are declared foreign to a GRF with respect to a pval, depending on the cluster maximum intensity value or their surface.

In this context, the parametric maps are described by the union of two classes of pixels: those representing a realization of a GRF modelling the normal case, and those that are foreign to the GRF. We expect these foreign elements to occur in regions with very high pixel intensity and/or in larger clusters taken at a specific threshold.

Modelling the parametric maps with GRF is only possible upon gaussianization (i.e. conversion of the empirical distributions into normal distributions) of the GRF marginal distributions. In practice, we only performed the gaussianization of the first-order marginal distribution of the GRF, i.e. the image intensity histogram, considering that the parameter maps under study are smooth enough. Therefore, the image intensity histogram was the only distribution to be gaussianized using an approach based on optimal transport (S1 Text). Thereby, the resulting histogram of intensity follows a normal distribution of zero mean with identical variance as the empirical native histogram.

To estimate the likelihood of a certain cluster to belong to a GRF termed G_r_, depending on the maximal intensity of this cluster, we used the following formulations taken from the theory of random fields (12). Considering x_0_ = t + H_0_ as the intensity peak of a cluster (at intensity threshold t), one can estimate the probability that a cluster having an intensity peak equal to x_0_, denoted 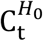 belongs to a realization of GRF. This probability can be seen as the likelihood of a cluster (taken at threshold t) of having an intensity peak higher or equal to t + H_0_:

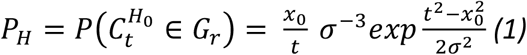

Furthermore, the approximation for the probability of a given cluster having a spatial extent S greater than S_0_ is given by the following formulation:

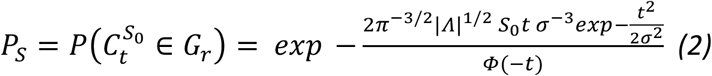

where

- Λ is the covariance matrix of the partial derivatives of G_r_
- *σ* is the standard deviation of G_r_
- Φ(−t) is the complementary cumulative distribution function

The comparison of these probability values against a pval (e.g. 0.05) was necessary to guide the decision regarding foreign cluster detection (i.e. a cluster is counted as detected if each probability is smaller or equal to pval).

This methodology (see Materials and Methods) was applied for a quantitative and qualitative analysis of fiber length and pore orientation isotropy differences, across all four FN variant networks in normal and tumor-like states. We were interested in determining whether the SPM analysis of the selected spatial fiber features could reveal significant variant-specific differences between the FN variant networks in normal (N) and tumor-like (T) states, among the different FN variant networks in a normal state, and finally, among the variant networks in a tumor-like state.

To set up this framework, we divided the available sets of confocal images (1024 × 1024 pixels, 0.27 9:/pixel; 70 images/variant for normal FN (N) and 65 images/variant for tumor-like FN networks) as follows. (1) For comparison of (N) vs (T) FN networks, we considered 50 (N) samples as the learning dataset, 20 (N) as test set for normal, and 65 (T) as test set for tumor-like networks. (2) Upon comparing (N) FN variant networks amongst each other, e.g. N_i_ vs N_j_, where i,j ∈ {B-A, B-A+, B+A-, B+A+}: 50 samples out of N_i_ were taken as the learning dataset, 20 out of N_i_ as test set, while 70 out of N_j_ comprised the test set. (3) Finally, we compared (T) FN variant meshworks amongst themselves, e.g. T_i_ vs T_j_, 50 samples out of T_i_ were taken as the learning dataset, 15 out of T_i_ as test set, while 65 out of T_j_ comprised the test set. In all scenarios, a cluster is detected as foreign to GRF if either *P*_*S*_ or *P*_*H*_ are less or equal than pval, pval ≤0.05. Anomalies in fiber length (Fig 5) and pore isotropy maps (Fig 6) were detected at a few intensity thresholds (e.g. 70,80,90) and (10,12,14), respectively (Fig 5, Fig 6). Thus, using our approach, differences in fiber length and pore orientation isotropy could be localized in regions formed at different intensity thresholds. This property is very useful to obtain a qualitative analysis of parametric maps, where clusters that are statistically different from a normal model can be localized.

**Fig 5.**
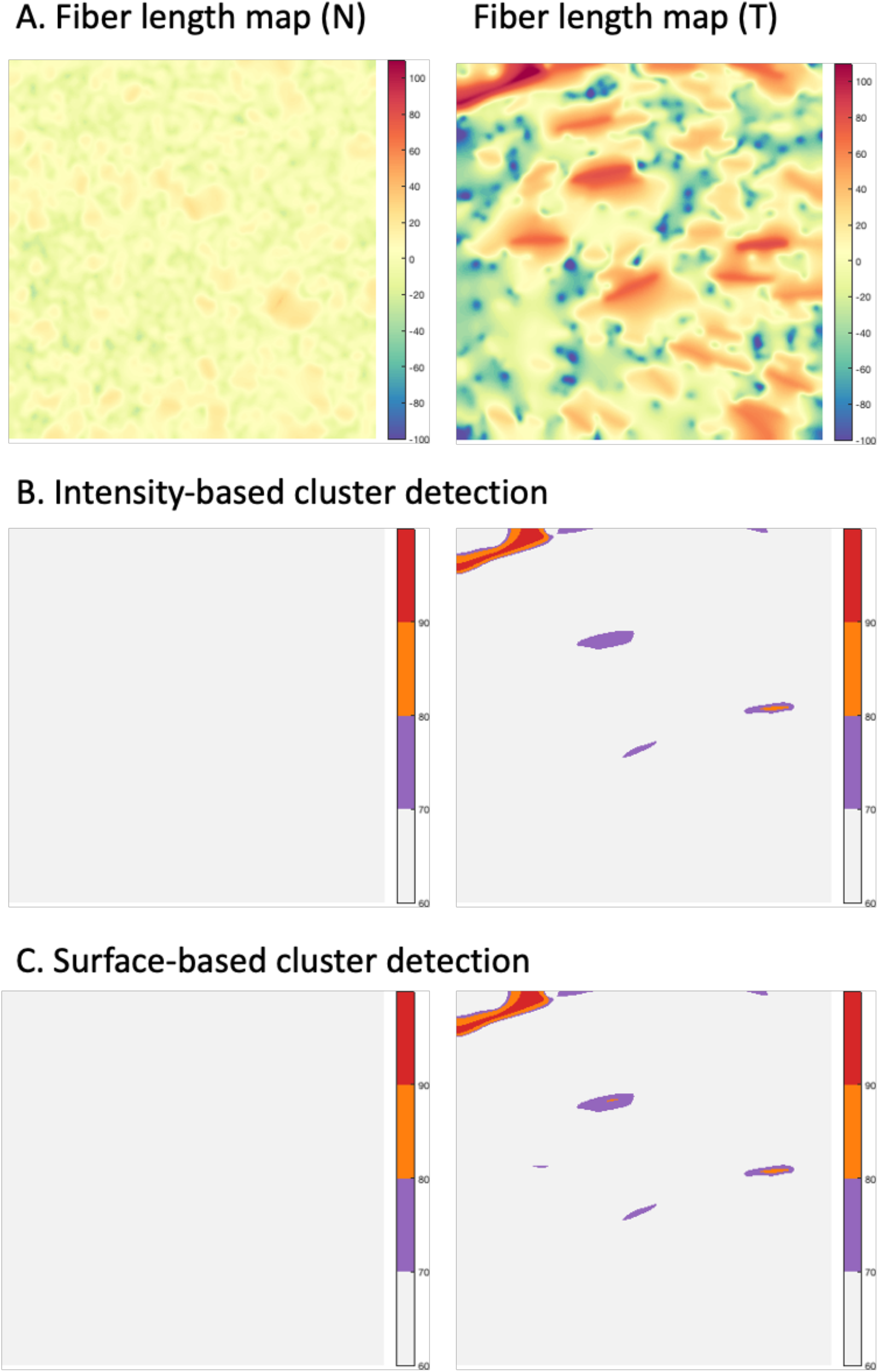
Qualitative analysis - Foreign cluster detection (with respect to the normal statistical model), applied to two samples of fiber length map (FN B-A+ tumor-like), 1024×1024 pixels (**A**). (**B**) and (**C**) depict the detected clusters (pval ≤0.05) at various intensity thresholds (70,80,90).

**Fig 6.**
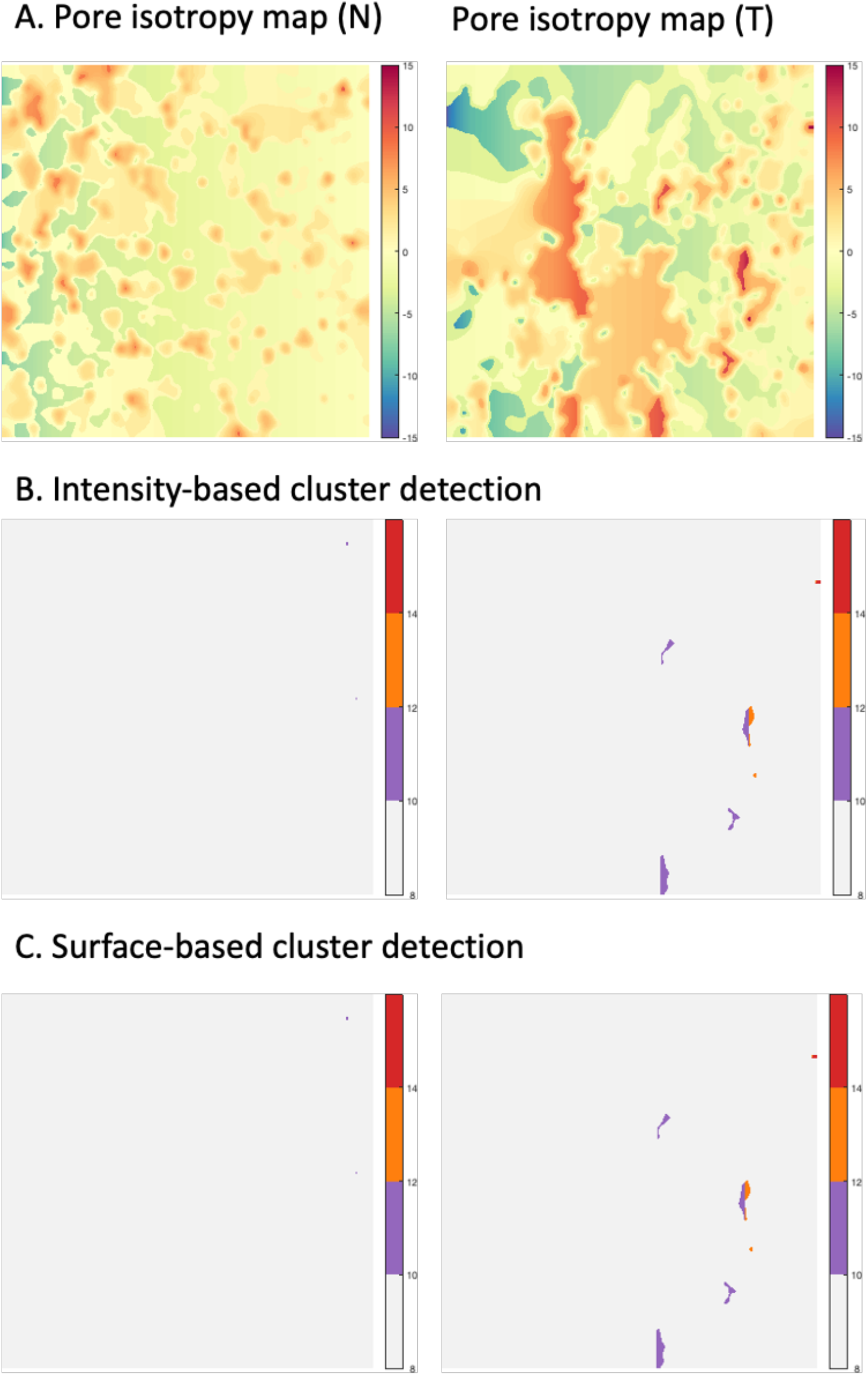
Qualitative analysis - Foreign cluster detection (with respect to the normal statistical model), applied to two samples of pore orientation isotropy map (FN B-A+ tumor-like), 1024×1024 pixels (**A**). (**B**) and (**C**) depict the detected clusters (pval ≤0.05) at various intensity thresholds (10,12,14).

For the quantitative analysis, we were interested in determining whether we could detect significant differences between FN variant networks through the average number of identified foreign clusters, as well as the average cluster area per image. It is noteworthy that the group for which clusters foreign to the GRF were found at superior thresholds, had higher parametric values than in the normal model. For example, if significantly different regions occur in the tumor-like matrices compared to the normal ones, with respect to fiber length, then fibers are statistically more elongated within tumor-like networks than normal counterparts. As shown in Fig 7, we found both fiber length and pore isotropy to be significantly different for pairwise comparisons of normal and tumor-like FN variant networks. Essentially, FN architecture in the tumor-like ECM, relative to normal ECM, is represented by statistically longer fibers (Fig 7A, bottom) with a more pronounced pore isotropy (Fig 7B, bottom). Detailed results, including the average area and number of identified foreign clusters to a GRF, are presented in S1 Table1-Table11 (S1 Text).

**Fig 7.**
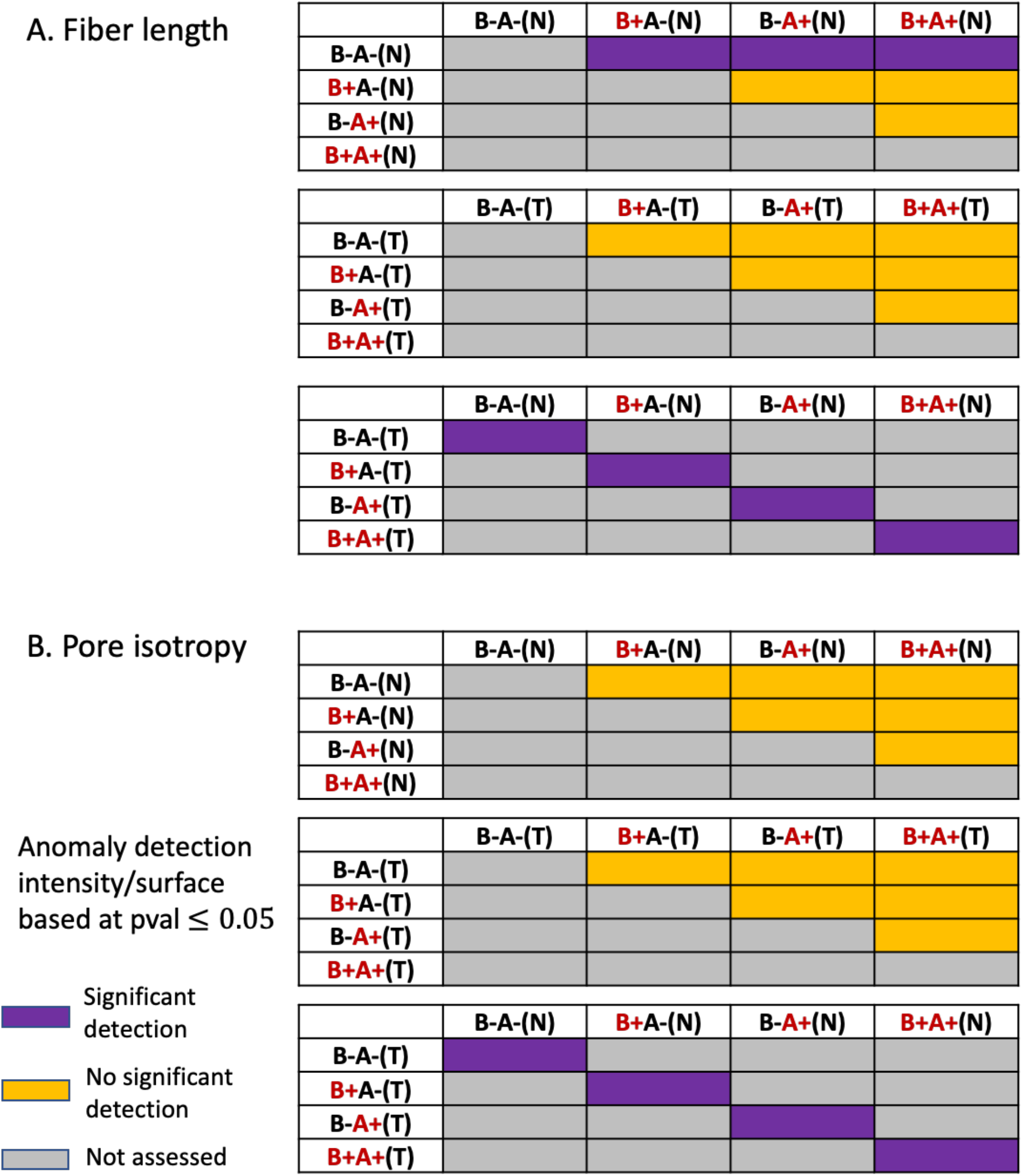
Quantitative analysis for detection of differences in fiber length and pore isotropy - Foreign cluster detection (with respect to the normal statistical model), applied for comparison of normal (N) and tumor-like (T) FN (1024×1024 pixels), where each matrix cell corresponds to a pairwise comparison between any two FN-specific variants. Detection is considered significant when at pval ≤ 0.05, the average number of foreign clusters per test database for one of the variants is higher than for the normal model): (**A**) Significant foreign cluster detection in fiber length maps. (**B**) Significant foreign cluster detection for pore orientation isotropy

The presence of FN Extra Domains (EDA and/or EDB) impacts FN assembly and fibrillar network organization by normal fibroblasts (9). This was demonstrated by local analysis of graph-based features (i.e. local node degree ratio, fiber thickness weighed by density, etc). An additional classification architecture was able to discriminate the four variant networks, on smaller images (512×512 pixels) with a relatively high accuracy, 83%. In accordance with these results, our current analysis of discriminating patterns across variant-specific FN fibrils showed that the inclusion of EDA and/or EDB in the cFN molecule, compared to pFN lacking these domains (B-A), affects the length of fibers assembled by normal (non-stimulated) fibroblasts (Fig 7A, top). Following activation, no significant differences in fiber length were observed across FN variant networks (Fig 7A, middle). This finding is not surprising, as activated fibroblasts in the tumor-like state are characterized by their enhanced contractility and elongated phenotype.

Interestingly, pore isotropy of the FN networks was similar for all four variants, in both normal and tumor-like states (Fig 7B, top and middle), suggesting that this parameter is not discriminating for the FN variants or activation state of the cells that assembled them. Intriguingly, our analyses revealed that pores of tumor-like matrices displayed a statistically significant orientation isotropy than matrices in the normal state (Fig 7B, bottom). Mechanisms underpinning this increase in random orientation of pores in the tumor-like FN networks remain to be elucidated.

## Conclusions

The proposed methodology was designed for the quantification of the differences in the spatial organization of ECM networks, in normal and tumor-like architectures. In this case, we chose to analyze the matrix patterns of four alternatively spliced FN variants, which we have previously been able to discriminate with different learning approaches and relying on graph-based feature analysis (9). The current computational pipeline is now available to be tested with a MATLAB GUI. Existing ECM analysis methods exploit alternative fiber detection techniques, such as Fast Fourier Transform bandpass filters (13), ridge detection (14), or fast discrete curvelet transform, which is arguably the best suited method to detect curvilinear anisotropic objects, among the previous options. The latter option was used in conjunction with a fiber extraction algorithm (6). Our method relies on a more flexible detection scheme using Gabor filters, thereby avoiding translation/rotation errors, and unlike other methods, associates graph networks to fiber morphological skeletons, enabling diverse fiber analyses. One such analysis, as seen in this work, provides a statistical assessment of the regions which are significantly different in terms of various geometric features.

The statistical approach can be evaluated on parametric maps at different thresholds, both quantitatively and qualitatively, whilst producing results that are more reliable and statistically relevant (by providing pval) than a simple hard thresholding of the parametric maps. The current analysis compares the fiber length and pore orientation isotropy parametric maps, of FN networks in normal and tumor-like states. Parametric maps can be extended to include other fiber or pore-specific parameters (e.g. fiber density, width, length, orientation, waviness, and straightness) useful for differentiating among various biological networks.

Computational analyses of ECM structures can provide essential information about their role in shaping the cellular microenvironment topology in health and during disease development. Prognostic ECM-specific signatures have already been inferred in cancer-related studies, diabetes, wound healing, and nerve repair. There is also a growing interest in the integration of cell and ECM analyses in a spatially resolved manner to further understand the interactions between cells and their matrix microenvironment (15). Our study proposes a versatile pipeline for the analysis of ECM organization in human tissue, in which a future integration of staining for cellular components can be envisioned. For instance, a combined local analysis of parametric maps and metrics describing the organization/morphology of adjacent cells could potentially help elucidate the complex interplay between cellular and non-cellular components.

## Materials and methods

### Materials and FN preparations

Recombinant human TGF-β1 was from R&D Systems Inc. (Minneapolis, MN, USA). All other chemicals and reagents were purchased from Sigma Aldrich (St Louis, MO, USA) unless otherwise stated. Purified recombinant FN variants were produced as previously described (9).

### Cells and culture conditions

*Fn1 -/-* mouse kidney fibroblasts (MFs) were generated and cultured as previously described (9). For experiments, FN was depleted from FCS using gelatin sepharose-4B columns (GE Healthcare, Uppsala, Sweden), and the culture medium was supplemented with Penicillin-Streptomycin 100 U/ml and TGF-β1 (5 ng/ml) where indicated. Absence of Mycoplasma sp. contamination was routinely verified by PCR as described elsewhere (16).

### Generation of MF-derived matrices, immunofluorescence staining and microscopy

MF-derived matrices were generated as described previously. For FN immunostaining, primary antibody (rabbit polyclonal anti-FN) was from MerckMillipore (Darmstadt, Germany). Fluorescently labeled (Alexa Fluor 488-conjugated) secondary antibody was from Thermo Fisher Scientific. After staining, the coverslips were mounted in ProLong® Gold antifade reagent (Thermo Fischer Scientific). Confocal imaging was performed on a Zeiss LSM710 confocal system equipped with a 10X/0.45 NA objective. For visual representation, image treatment was performed using Fiji (17).

### Statistical analysis based on GRF

Statistical parametric maps (SPMs) are used to evaluate the probability of change in every pixel by using decision tests based on the magnitude of the SPM values (i.e. the peak intensity of a cluster in SPM) and on the spatial extent of these clusters formed at a certain threshold. In our experiments, we focused on two different parametric maps representing the length of fibers and the isotropy of pore angles. We embedded the SPM framework, initially developed to analyze single datasets independently, into a machine learning paradigm. To evaluate the differences between two given groups of maps (e.g., FN variants in normal state vs corresponding FN variants tumor-like state), we considered one of the groups as the normal realization of the GRF which we divide into a learning set and a smaller test set. The second group is tested for foreign regions, therefore all the images belonging to this group are considered part of the test set. The proposed method learns the normal GRF model specific parameters, i.e. the average value of Λ_*j*_, *σ*_*j*_, from the training set. These learnt parameters are subsequently used to compute the two relevant probabilities, P_H_ and P_S_.

In the learning phase, we learn the model parameter, as follows.

For all images *I*_*j*_ (previously gaussianized (S1 Fig)) in the learning set:

- Compute *Λ*_j_ (empirical estimator of the covariance of partial derivatives of *I*_*j*_). If for an image function *f ϵ R*^2^, we consider its gradient vector ∇*f* = (*f*_*x*_, *f*_*y*_) = (*∂ f / ∂ x, ∂ f / ∂ y*) then Λ_*j*_ = *cov* (*f*_*x*_, *f*_*y*_)= *E*[(*f*_*x*_ − *E*[*f*_*x*_]) (*f*_*y*_ – *E*[*f*_*y*_])].
- Compute *σ*_*j*_, as the *I*_*j*_ ‘s sample standard deviation.

The last step involves storing the average Λ_*m*_, *σ*_*m*_ of the learning dataset.

The test phase is where the decision whether the clusters, taken at a certain threshold t, are considered foreign to the GRF, is made, according to a given pval.

For all the images *I*_*j*_ in the test set:

- Gaussianize *I*_*j*_ so that it can be approximated by a realization of a GRF. The result is a new image *I*_*g*_, whose histogram is Gaussian with identical variance to that of *I*_*j*_.
- For a given list of thresholds *T* = (*t*_1_, *t*_2_, …, *t*_*n*_):
  ▪ Binarize *I*_*g*_ according to the threshold *t*_*i*_
  ▪ Find the list of connected components in the binary image resulted from thresholding and for every (labeled) connected component (*l*_1_, *l*_2_, …, *l*_*p*_):
    - Compute *P*_*H*_ (Eq.1), using the learnt model parameter *σ*_*m*_. If its value is lower than a chosen pval, the cluster is considered a foreign element to a GRF.
    - Compute *P*_*S*_ (Eq.2), using the learnt model parameters Λ_*m*_, *σ*_*m*_. If its value is lower than a chosen pval, the cluster is considered a foreign element to a GRF.

### Test for anomaly detection in a GRF using simulation

To better illustrate the principle behind the proposed approach to detect statistical abnormalities in a GRF realization, we considered a simple scenario which includes a synthetic example (normal), representing a simulation of a GRF of zero mean. To this example we added 6 foreign objects (abnormal), representing ellipses with different surfaces, and intensity levels. The aim of this example is to showcase the suitability of our tool when detecting anomalies at various thresholds, based on the maximum intensity of the different regions detected at each threshold, or their spatial extent. Specifically, in this test, we were interested in verifying whether our approach can detect the foreign ellipses, at various thresholds, for pval ≤0.05 based on either an intensity or a surface criterion.

Our method learns the GRF model parameters from the normal example and then uses these to compute the two probabilities of belonging to GRF for each region at various thresholds in the abnormal example. As shown in Fig 8, the current method, compared to a naïve hard thresholding (threshold equal to 5), is better suited to detect only the abnormal elements, i.e. ellipses, thresholds equal to (5,10,15,20). A possible decision could jointly consider the two criteria, i.e. intensity and surface-based detection, and thus lead to the selection of the 6 ellipses together with a few false positive alarms.

**Fig 8.**
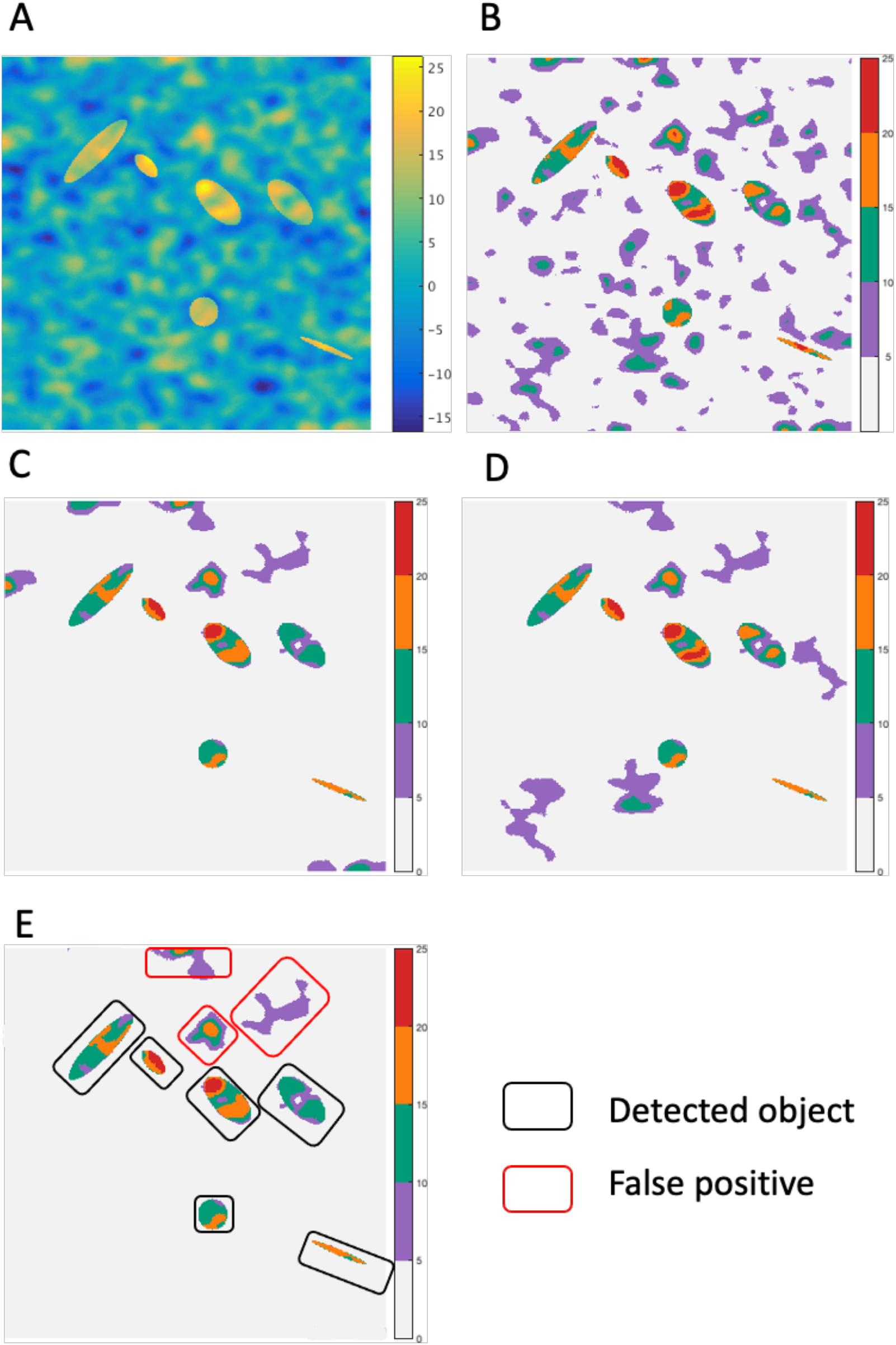
(**A**) Realization of a GRF of zero-mean and addition of 6 different sized ellipses. (**B**) Hard thresholding (t = 5 pixels). (**C**) Intensity-based detection at pval ≤0.05, at thresholds equal to (5,10,15,20, see colorbar). (**D**) Surface-based detection at thresholds equal to (5,10,15,25, see colorbar). (**E**) Example of joint decision of cluster selection based on intensity and surface criteria

## Acknowledgements

We thank the members of the Adhesion Signaling and Stromal Reprogramming in the Tumor Microenvironment team for critical discussion on the manuscript, iBV PRISM imaging platform for the use of their machines and support, in particular, Sébastien Schaub.

## Competing interests

The authors declare no competing interests.

## Supporting Information (S1)

### S1 text: Supporting Methods

#### Graph-based representation on top of Gabor filters

We defined a set of Gabor kernels *g*_*k*_, characterized by an exponential term which provides the shape of a bivariate gaussian kernel multiplied by a cosine function which describes its oscillations in space:

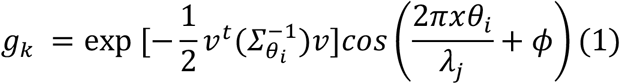

where we denote by *ν* = (*x, y*)^*t*^, the 2D coordinate vector, indicating pixels localisation in a bidimendional Cartesian coordinate system.

To capture the various geometrical properties of the FN fibers using Gabor filters, we have constructed a set of Gabor kernels *g*_*k*_, *k* ∈ {1,2, … 60}, defined by the following parameters:

- Fiber orientation is represented by *θ*_*i*_, *θ*_*i*_ = *iπ*/20 where D = {0,1,2, … 19}.
- Fiber thickness is denoted as *λ*_*j*_, *j* = {1,2,3} and corresponds to the wavelength (in pixels) of the cosine term, the values of which are equal to *λ*_*j*_/2 and vary between 3 and 5 pixels. Therefore, the thinnest fibers are detected when *λ*_*j*_= 6 pixels, medium thickness fibers correspond to *λ*_*j*_= 8 pixels, while the thickest are characterized by *λ*_*j*_ = 10 pixels.
- For accurate localization of fibers, the phase of the cosine function, ϕ, is set to {0}.
- The spatial support of the kernels is given by *σ*_*x*_ = 5 pixels and *σ*_*y*_ = 3 pixels, indicating an anisotropic filter that is appropriate for fiber detection.

At any given location within the filtered image sample, the Gabor kernel that returns the highest coefficient modulus was retained. Once fibers were enhanced using Gabor filters, we extracted their morphological skeleton, using different morphological operations (e.g. binarization with hysteresis, morphological thinning, etc.). The graph-based representation of the fiber skeletons was subsequently derived, using a toolbox that produces a network graph assigned to a morphological skeleton (1). To reconstruct the missing fibers, we used Dijkstra’s weighted shortest path algorithm, which finds for each node (i.e. fiber end), the shortest path to other extremities or nodes of the skeleton graph, which are within a given distance to the considered node. Among all possible paths, we chose the minimal one, relying on the intensity from the Gabor filtered image. Additionally, the reconnection of the fibers was made using the angle of the Gabor maximum response as a guideline, which resulted in the reconnection of the fibers within a predefined cone sector around this local fiber orientation.

### Parametric maps derivation

To build a fiber length map, we relied on the simplified graph-based representation, where we kept the location of the nodes (fiber ends and crosslinks) and replaced the length of the fiber with a connecting line. During the next step, we identified the 2D pixels coordinates that approximate the straight line between the nodes, (using Bresenham’s line algorithm) and assigned the length value of the connecting line to these pixels. The last step for generating the fiber length map, included the extrapolation of the fiber length values (2) and smoothing of this result with a Gaussian kernel.

Concerning the pore isotropy parametric maps, the starting point was the skeleton graph. Pore orientation was computed by first fitting ellipses to each pore and was subsequently obtained by measuring the angle *θ*_*i*_ between the horizontal axis and the major ellipse axis. Pore median value is then subtracted from each *θ*_*i*_. Finally, the inverse absolute value of the resulting individual score per pore (indicative of the orientation isotropy) will be assigned to all pixels filling its corresponding surface, and subsequently smoothed out with a Gaussian kernel.

### 1D Optimal Transport

The approach that was considered for map gaussianization was based on the optimal transport framework (3). More specifically, the problem of “converting” the empirical intensity distribution of the fiber length map into a normal distribution with the same variance, can be approached by performing the 1D optimal transport between the two distributions. Briefly, for a map I with an empirical mean and variance of intensity, we consider a second image J, whose intensity pixels follow a normal distribution (S1 Fig). The 1D optimal transport problem will determine how to optimally permute the pixels indices in J, to “recreate” the image I. Since the intensity of the pixels in the permuted version of J does not change, the result will resemble I, but will have the histogram of J (4).

### Gaussian Random Fields

Given a complete probability space (Ω, F, P) and T a topological space, then a random field of real values is a measurable application *X*: Ω → *R*^*T*^. The finite-dimensional (*F*_*d*_) distributions of X are defined as the set of functions 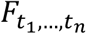, where 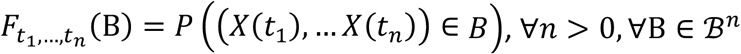, F(B) being the Borel sigma algebra on Q.

A particular class of random fields is represented by the Gaussian random fields, whose marginal (*F*_*d*_) distributions are Gaussian vectors *X* = (*X* _(1)_, …, *X* _(*n*)_), characterized by the probability density function (5):

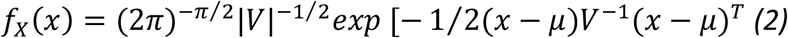

**S1 Fig.**
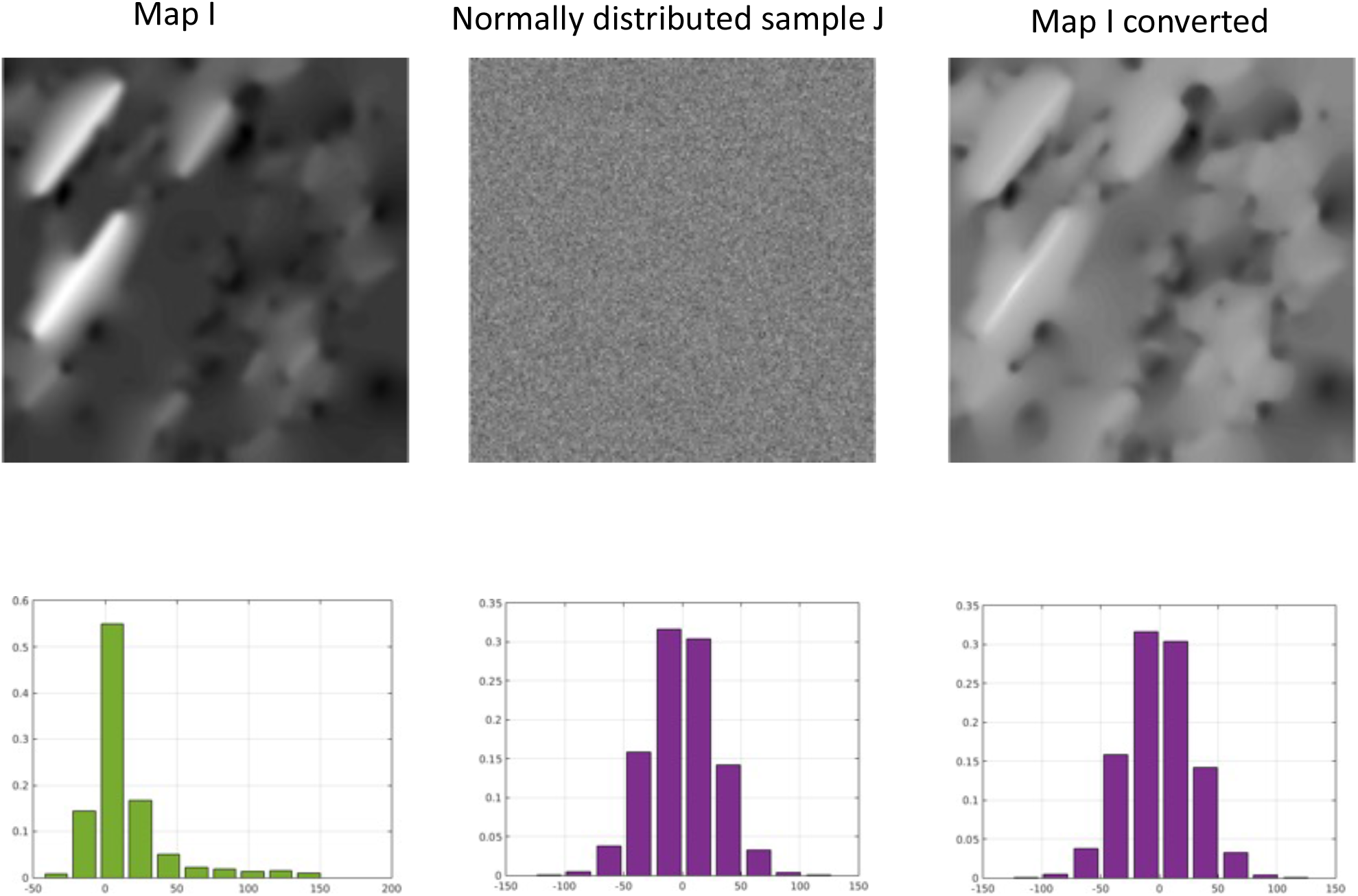
Gaussianization through optimal transport: First row illustrates an example of a fiber length parametric map (Map I), a generated image J with normally distributed intensity around 0, and equal variance to that of Map I, and, finally, the converted map I with a corresponding normally distributed intensity. Second row shows the corresponding intensity histograms

Where 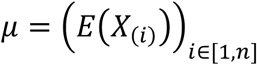 is the expectation and 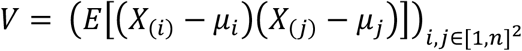 is the covariance matrix.

The expectation (or mean value) of the number of clusters that appear at a threshold t, of an image modelled by a zero-mean, homogeneous Gaussian field of dimension 2, is the following (5):

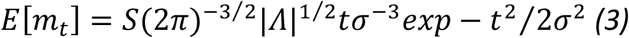

where:

- *m*_*t*_ represents the number of clusters at a certain threshold t
- S is the number of pixels of the image
- Λ is the covariance matrix of partial derivatives of the GRF
- *σ* is the standard deviation of the GRF

Similarly, the mean value of the number of clusters at a threshold t + l_9_ can be written as such:

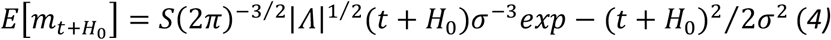

Considering *x*_0_ = *t* + *H*_0_ as the intensity peak of a cluster (at threshold t), one can estimate the probability that a cluster (at a threshold t, having an intensity peak equal to *x*_0_, denoted 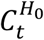) belongs to a realization of this GRF. This probability can be seen as the likelihood of a cluster (taken at threshold t) of having an intensity peak higher or equal to *t* + *H*_0_:

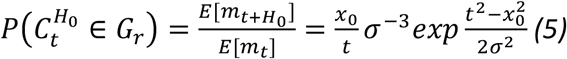

Next, we are interested in the estimation of the probability that a cluster (at a threshold t) belongs to a realization of GRF, depending on its surface (spatial extent - number of pixels). To estimate the number of pixels (*n*_*t*_) of a cluster at a threshold t, we use the following equation (6):

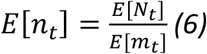

where *N*_*t*_ is the number of pixels of higher intensity than t, and *m*_*t*_ is the number of clusters at threshold t. Since the intensity values follow a normal (zero mean value) distribution, the expectation of *N*_*t*_ is the following:

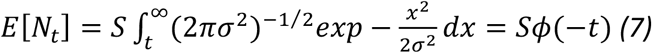

where *ϕ* (−*t*) is the complementary cumulative distribution function. It follows then (Eq. 3, 6, 7) that one can approximate the mean value of *n*_*t*_, accordingly:

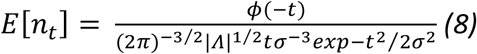

Furthermore, it has been shown in (7) that *n*_*t*_ follows an exponential distribution law, which is commonly defined by a parameter *λ*_*t*_, (the inverse of the mean expected value of the random variable).

Consequently

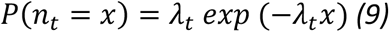

*where*

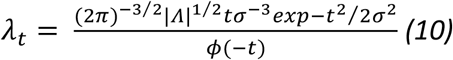

It follows then that the approximation for the probability of a given cluster having a spatial extent S greater than *S*_0_ is given by the following formulation:

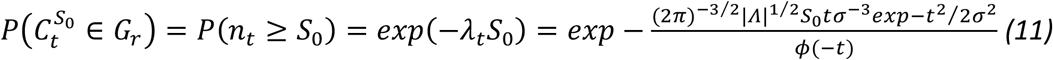

### Quantitative results (area and number) of detected clusters foreign to a GRF

**S1 Table 1.** Foreign cluster detection (with respect to the normal statistical model), applied for the comparison of B-A- and B-A+ FN (1024×1024 pixels) fiber length maps. Clusters are detected according to intensity and surface criteria, at pval ≤0.05, for various thresholds 70,80,90. The average number (count) and area of detected clusters are recorded, where the results corresponding to the normal model are shown in the first half of the table (B-A vs B-A).

|  | B-A- vs B-A- | Thresholds |  |  |  | B-A- vs B-A+ | Thresholds |  |  |
| --- | --- | --- | --- | --- | --- | --- | --- | --- | --- |
|  |  | 70 | 80 | 90 |  |  | 70 | 80 | 90 |
| Intensity | Count | 0.10 | 0.00 | 0.00 | Intensity | Count | 0.29 | 0.11 | 0.03 |
|  | Area | 79.00 | 0.00 | 0.00 |  | Area | 159.95 | 48.00 | 18.00 |
| Surface | Count | 0.15 | 0.10 | 0.00 | Surface | Count | 0.63 | 0.37 | 0.17 |
|  | Area | 44.33 | 4.00 | 0.00 |  | Area | 0.63 | 0.37 | 0.17 |

**S1 Table 2.** Foreign cluster detection (with respect to the normal statistical model), applied for the comparison of B-A- and B+A-FN (1024×1024 pixels) fiber length maps. Clusters are detected according to intensity and surface criteria, at pval ≤0.05, for various thresholds 70,80,90. The average number (count) and area of detected clusters are recorded, where the results corresponding to the normal model are shown in the first half of the table (B-A vs B-A).

|  | B-A- vs B-A- | Thresholds |  |  |  | B-A- vs B+A- | Thresholds |  |  |
| --- | --- | --- | --- | --- | --- | --- | --- | --- | --- |
|  |  | 70 | 80 | 90 |  |  | 70 | 80 | 90 |
| Intensity | Count | 0.00 | 0.00 | 0.00 | Intensity | Count | 0.19 | 0.04 | 0.00 |
|  | Area | 0.00 | 0.00 | 0.00 |  | Area | 228.85 | 89.00 | 0.00 |
| Surface | Count | 0.07 | 0.00 | 0.00 | Surface | Count | 0.59 | 0.34 | 0.09 |
|  | Area | 3.00 | 0.00 | 0.00 |  | Area | 85.20 | 19.92 | 9.00 |

**S1 Table 3.** Foreign cluster detection (with respect to the normal statistical model), applied for the comparison of B-A- and B+A+ FN (1024×1024 pixels) fiber length maps. Clusters are detected according to intensity and surface criteria, at pval ≤0.05, for various thresholds 70,80,90. The average number (count) and area of detected clusters are recorded, where the results corresponding to the normal model are shown in the first half of the table (B-A vs B-A).

|  | B-A- vs B-A- | Thresholds |  |  |  | B-A- vs B+A+ | Thresholds |  |  |
| --- | --- | --- | --- | --- | --- | --- | --- | --- | --- |
|  |  | 70 | 80 | 90 |  |  | 70 | 80 | 90 |
| Intensity | Count | 0.00 | 0.00 | 0.00 | Intensity | Count | 0.00 | 0.00 | 0.00 |
|  | Area | 0.00 | 0.00 | 0.00 |  | Area | 0.00 | 0.00 | 0.00 |
| Surface | Count | 0.1 | 0.00 | 0.00 | Surface | Count | 0.77 | 0.39 | 0.16 |
|  | Area | 17.5 | 0.00 | 0.00 |  | Area | 63.65 | 18.44 | 6.18 |

**S1 Table 4.**
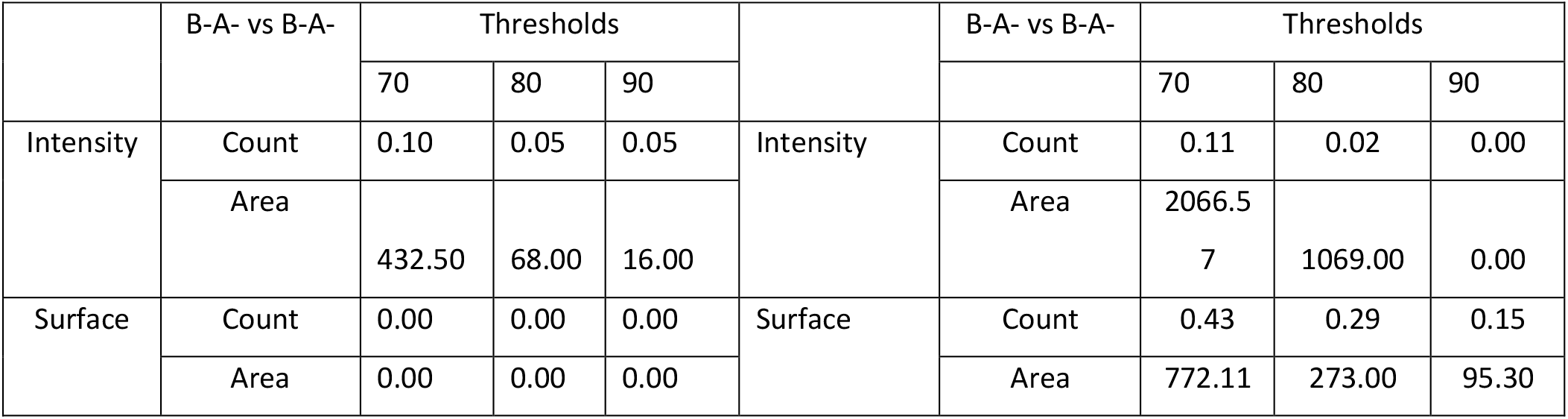
Foreign cluster detection (with respect to the normal statistical model), applied for the comparison of B-A- (N) and B-A- (T) FN (1024×1024 pixels) fiber length maps. Clusters are detected according to intensity and surface criteria, at pval ≤0.05, for various thresholds 70,80,90. The average number (count) and area of detected clusters are recorded, where the results corresponding to the normal model are shown in the first half of the table.

|  | B-A- vs B-A- | Thresholds |  |  |  | B-A- vs B-A- | Thresholds |  |  |
| --- | --- | --- | --- | --- | --- | --- | --- | --- | --- |
|  |  | 70 | 80 | 90 |  |  | 70 | 80 | 90 |
| Intensity | Count | 0.10 | 0.05 | 0.05 | Intensity | Count | 0.11 | 0.02 | 0.00 |
|  | Area | 432.50 | 68.00 | 16.00 |  | Area | 2066.5<br>7 | 1069.00 | 0.00 |
| Surface | Count | 0.00 | 0.00 | 0.00 | Surface | Count | 0.43 | 0.29 | 0.15 |
|  | Area | 0.00 | 0.00 | 0.00 |  | Area | 772.11 | 273.00 | 95.30 |

**S1 Table 5.** Foreign cluster detection (with respect to the normal statistical model), applied for the comparison of B+A- (N) and B+A- (T) FN (1024×1024 pixels) fiber length maps. Clusters are detected according to intensity and surface criteria, at pval ≤0.05, for various thresholds 70,80,90. The average number (count) and area of detected clusters are recorded, where the results corresponding to the normal model are shown in the first half of the table.

|  | B+A- vs B+A- | Thresholds |  |  |  | B+A- vs B+A- | Thresholds |  |  |
| --- | --- | --- | --- | --- | --- | --- | --- | --- | --- |
|  |  | 70 | 80 | 90 |  |  | 70 | 80 | 90 |
| Intensity | Count | 0.00 | 0.00 | 0.00 | Intensity | Count | 0.88 | 0.51 | 0.26 |
|  | Area | 0.00 | 0.00 | 0.00 |  | Area | 3608.82 | 2194.45 | 1165.65 |
| Surface | Count | 0.15 | 0.10 | 0.10 | Surface | Count | 0.92 | 0.89 | 0.78 |
|  | Area | 220.67 | 44.50 | 7.00 |  | Area | 3127.58 | 1315.53 | 506.22 |

**S1 Table 6.** Foreign cluster detection (with respect to the normal statistical model), applied for the comparison of B-A+ (N) and B-A+ (T) FN (1024×1024 pixels) fiber length maps. Clusters are detected according to intensity and surface criteria, at pval ≤0.05, for various thresholds 70,80,90. The average number (count) and area of detected clusters are recorded, where the results corresponding to the normal model are shown in the first half of the table.

|  | B-A+ vs B-A+ | Thresholds |  |  |  | B-A+ vs B-A+ | Thresholds |  |  |
| --- | --- | --- | --- | --- | --- | --- | --- | --- | --- |
|  |  | 70 | 80 | 90 |  |  | 70 | 80 | 90 |
| Intensity | Count | 0.00 | 0.00 | 0.00 | Intensity | Count | 0.72 | 0.40 | 0.23 |
|  | Area | 0.00 | 0.00 | 0.00 |  | Area | 6038.85 | 6265.31 | 6553.13 |
| Surface | Count | 0.15 | 0.10 | 0.10 | Surface | Count | 0.78 | 0.54 | 0.42 |
|  | Area | 38.00 | 5.00 | 1.50 |  | Area | 5337.84 | 4563.03 | 3591.33 |

**S1 Table 7.** Foreign cluster detection (with respect to the normal statistical model), applied for the comparison of B+A+ (N) and B+A+ (T) FN (1024×1024 pixels) fiber length maps. Clusters are detected according to intensity and surface criteria, at pval ≤0.05, for various thresholds 70,80,90. The average number (count) and area of detected clusters are recorded, where the results corresponding to the normal model are shown in the first half of the table.

|  | B+A+ vs B+A+ | Thresholds |  |  |  | B+A+ vs B+A+ | Thresholds |  |  |
| --- | --- | --- | --- | --- | --- | --- | --- | --- | --- |
|  |  | 70 | 80 | 90 |  |  | 70 | 80 | 90 |
| Intensity | Count | 0.00 | 0.00 | 0.00 | Intensity | Count | 0.94 | 0.49 | 0.29 |
|  | Area | 0.00 | 0.00 | 0.00 |  | Area | 3609.21 | 2509.34 | 1511.63 |
| Surface | Count | 0.05 | 0.00 | 0.00 | Surface | Count | 0.94 | 0.74 | 0.71 |
|  | Area | 17.00 | 0.00 | 0.00 |  | Area | 3298.38 | 1574.79 | 675.59 |

**S1 Table 8.** Foreign cluster detection (with respect to the normal statistical model), applied for the comparison of B-A- (N) and B-A- (T) FN (1024×1024 pixels) pore isotropy maps. Clusters are detected according to intensity and surface criteria, at pval ≤0.05, for various thresholds 10,12,14. The average number (count) and area of detected clusters are recorded, where the results corresponding to the normal model are shown in the first half of the table.

|  | B-A- vs B-A- | Thresholds |  |  |  | B-A- vs B-A- | Thresholds |  |  |
| --- | --- | --- | --- | --- | --- | --- | --- | --- | --- |
|  |  | 10 | 12 | 14 |  |  | 10 | 12 | 14 |
| Intensity | Count | 0.05 | 0.00 | 0.00 | Intensity | Count | 0.12 | 0.03 | 0.00 |
|  | Area | 1041.00 | 0.00 | 0.00 |  | Area | 1216.63 | 429.50 | 0.00 |
| Surface | Count | 0.25 | 0.20 | 0.00 | Surface | Count | 0.31 | 0.20 | 0.06 |
|  | Area | 370.20 | 37.00 | 0.00 |  | Area | 480.10 | 99.15 | 37.25 |

**S1 Table 9.** Foreign cluster detection (with respect to the normal statistical model), applied for the comparison of B+A- (N) and B+A- (T) FN (1024×1024 pixels) pore isotropy maps. Clusters are detected according to intensity and surface criteria, at pval ≤0.05, for various thresholds 10,12,14. The average number (count) and area of detected clusters are recorded, where the results corresponding to the normal model are shown in the first half of the table.

|  | B+A- vs B+A- | Thresholds |  |  |  | B+A- vs B+A- | Thresholds |  |  |
| --- | --- | --- | --- | --- | --- | --- | --- | --- | --- |
|  |  | 10 | 12 | 14 |  |  | 10 | 12 | 14 |
| Intensity | Count | 0.10 | 0.05 | 0.00 | Intensity | Count | 0.35 | 0.05 | 0.00 |
|  | Area | 947.50 | 292.00 | 0.00 |  | Area | 979.43 | 558.33 | 0.00 |
| Surface | Count | 0.30 | 0.05 | 0.00 | Surface | Count | 0.42 | 0.22 | 0.15 |
|  | Area | 400.83 | 278.00 | 0.00 |  | Area | 482.00 | 187.79 | 39.30 |

**S1 Table 10.** Foreign cluster detection (with respect to the normal statistical model), applied for the comparison of B-A+ (N) and B-A+ (T) FN (1024×1024 pixels) pore isotropy maps. Clusters are detected according to intensity and surface criteria, at pval ≤0.05, for various thresholds 10,12,14. The average number (count) and area of detected clusters are recorded, where the results corresponding to the normal model are shown in the first half of the table.

|  | B-A+ vs B-A+ | Thresholds |  |  |  | B-A+ vs B-A+ | Thresholds |  |  |
| --- | --- | --- | --- | --- | --- | --- | --- | --- | --- |
|  |  | 10 | 12 | 14 |  |  | 10 | 12 | 14 |
| Intensity | Count | 0.15 | 0.00 | 0.00 | Intensity | Count | 0.54 | 0.14 | 0.02 |
|  | Area | 561.67 | 0.00 | 0.00 |  | Area | 1169.49 | 513.44 | 238.00 |
| Surface | Count | 0.15 | 0.05 | 0.00 | Surface | Count | 0.52 | 0.34 | 0.15 |
|  | Area | 478.33 | 132.00 | 0.00 |  | Area | 651.68 | 162.27 | 43.50 |

**S1 Table 11.** Foreign cluster detection (with respect to the normal statistical model), applied for the comparison of B+A+ (N) and B+A+ (T) FN (1024×1024 pixels) pore isotropy maps. Clusters are detected according to intensity and surface criteria, at pval ≤0.05, for various thresholds 10,12,14. The average number (count) and area of detected clusters are recorded, where the results corresponding to the normal model are shown in the first half of the table.

|  | B+A+ vs B+A+ | Thresholds |  |  |  | B+A+ vs B+A+ | Thresholds |  |  |
| --- | --- | --- | --- | --- | --- | --- | --- | --- | --- |
|  |  | 10 | 12 | 14 |  |  | 10 | 12 | 14 |
| Intensity | Count | 0.15 | 0.05 | 0.05 | Intensity | Count | 0.11 | 0.03 | 0.00 |
|  | Area | 3933.33 | 2804.00 | 469.00 |  | Area | 1034.43 | 368.00 | 0.00 |
| Surface | Count | 0.35 | 0.20 | 0.05 | Surface | Count | 0.18 | 0.09 | 0.05 |
|  | Area | 1565.43 | 771.50 | 89.00 |  | Area | 633.42 | 170.33 | 35.33 |

## Notes

### Competing Interest Statement

The authors have declared no competing interest.

## References

1. Frantz C, Stewart KM, Weaver VM. The extracellular matrix at a glance. Journal of Cell Science. 2010 Dec 15;123(24):4195–200.

2. Afratis NA, Sagi I. Novel Approaches for Extracellular Matrix Targeting in Disease Treatment. In: Vigetti D, Theocharis AD, editors. The Extracellular Matrix [Internet]. New York, NY: Springer New York; 2019 [cited 2021 Oct 11]. p. 261–75. (Methods in Molecular Biology; vol. 1952). Available from: http://link.springer.com/10.1007/978-1-4939-9133-4_21

3. Nia HT, Munn LL, Jain RK. Physical traits of cancer. Science. 2020 Oct 30;370(6516):eaaz0868.

4. Conklin MW, Eickhoff JC, Riching KM, Pehlke CA, Eliceiri KW, Provenzano PP, et al. Aligned collagen is a prognostic signature for survival in human breast carcinoma. American Journal of Pathology. 2011;178(3):1221–32.

5. Provenzano PP, Eliceiri KW, Campbell JM, Inman DR, White JG, Keely PJ. Collagen reorganization at the tumor-stromal interface facilitates local invasion. BMC Med. 2006 Dec;4(1):38.

6. Burke K, Smid M, Dawes RP, Timmermans MA, Salzman P, van Deurzen CHM, et al. Using second harmonic generation to predict patient outcome in solid tumors. BMC Cancer. 2015 Dec;15(1):929.

7. Hynes RO. The extracellular matrix: Not just pretty fibrils. Science. 2009 Nov 27;326(5957):1216–9.

8. Barker TH, Engler AJ. The provisional matrix: setting the stage for tissue repair outcomes. Matrix Biology. 2017 Jul;60–61:1–4.

9. Efthymiou G, Radwanska A, Grapa AI, Beghelli-de la Forest Divonne S, Grall D, Schaub S, et al. Fibronectin Extra Domains tune cellular responses and confer topographically distinct features to fibril networks. Journal of Cell Science [Internet]. 2021 Feb 24 [cited 2021 May 31];134(jcs252957). Available from: https://doi.org/10.1242/jcs.252957

10. Efthymiou G, Saint A, Ruff M, Rekad Z, Ciais D, Van Obberghen-Schilling E. Shaping Up the Tumor Microenvironment With Cellular Fibronectin. Front Oncol. 2020 Apr 30;10:641.

11. Lafarge F, Descombes X, Zerubia J, Mathieu S. Détection de feux de forêt par analyse statistique d’évènements rares à partir d’images infrarouges thermiques Forest fire detection by statistical analysis of rare events from thermical infrared images.

12. Adler RJ. 5. Some Expectations. In: The Geometry of Random Fields [Internet]. Society for Industrial and Applied Mathematics; 2010 [cited 2021 Jun 2]. p. 93– 121. (Classics in Applied Mathematics). Available from: https://doi.org/10.1137/1.9780898718980.ch5

13. Morrill EE, Tulepbergenov AN, Stender CJ, Lamichhane R, Brown RJ, Lujan TJ. A validated software application to measure fiber organization in soft tissue. Biomech Model Mechanobiol. 2016 Dec;15(6):1467–78.

14. Wershof E, Park D, Barry DJ, Jenkins RP, Rullan A, Wilkins A, et al. A FIJI macro for quantifying pattern in extracellular matrix. Life Sci Alliance. 2021 Mar;4(3):e202000880.

15. Vasiukov G, Novitskaya T, Senosain MF, Camai A, Menshikh A, Massion P, et al. Integrated Cells and Collagen Fibers Spatial Image Analysis. Front Bioinform. 2021 Nov 8;1:758775.

16. Kong F, James G, Gordon S, Zelynski A, Gilbert GL. Species-Specific PCR for Identification of Common Contaminant Mollicutes in Cell Culture. Appl Environ Microbiol. 2001 Jul;67(7):3195–200.

17. Schindelin J, Arganda-Carreras I, Frise E, Kaynig V, Longair M, Pietzsch T, et al. Fiji: an open-source platform for biological-image analysis. Nat Methods. 2012 Jul;9(7):676–82.

18. Poline JB, Worsley KJ, Evans AC, Friston KJ. Combining Spatial Extent and Peak Intensity to Test for Activations in Functional Imaging. 1997.

## References

1. Kollmannsberger P, Kerschnitzki M, Repp F, Wagermaier W, Weinkamer R, Fratzl P. The small world of osteocytes: Connectomics of the lacuno-canalicular network in bone. New Journal of Physics. 2017 Jul 1;19(7).

2. D’Errico J. Interpolation and extrapolation of elements in a 2D array [Internet]. John D’Errico (2021). inpaint_nans (https://www.mathworks.com/matlabcentral/fileexchange/4551-inpaint_nans), MATLAB Central File Exchange. Retrieved October 12, 2021.; Available from: https://www.mathworks.com/matlabcentral/fileexchange/4551-inpaint_nans

3. Peyré G, Cuturi M. Computational Optimal Transport. arXiv:180300567 [stat] [Internet]. 2020 Mar 18 [cited 2021 Oct 12]; Available from: http://arxiv.org/abs/1803.00567

4. Peyré G. The Numerical Tours of Signal Processing - Advanced Computational Signal and Image Processing. [Internet]. IEEE Computing in Science and Engineering; 2011. Available from: https://hal.archives-ouvertes.fr/hal-00519521/document

5. Adler RJ. 5. Some Expectations. In: The Geometry of Random Fields [Internet]. Society for Industrial and Applied Mathematics; 2010 [cited 2021 Jun 2]. p. 93–121. (Classics in Applied Mathematics). Available from: https://doi.org/10.1137/1.9780898718980.ch5

6. Friston K, Worsley K, Frackowiak R, Mazziotta J, Evans A. Assessing the Significance of Focal Activations Using Their Spatial Extent. Vol. 1, + Human Brain Mapping. 1994 p. 210–20.

7. Belyaev Yu K, Nosko VP. Characteristics of Excursions above a High Level for a Gaussian Process and Its Envelope. Theory Probab Appl. 1969 Jan;14(2):296–309.

